# UNLOCKING ROBOTIC POTENTIAL THROUGH MODERN ORGAN SEGMENTATION

**DOI:** 10.1101/2023.11.18.567632

**Authors:** Ansh Chaudhary

## Abstract

Deep learning has revolutionized the approach to complex data-driven problems, specifically in medical imaging, where its techniques have significantly raised efficiency in organ segmentation. The urgent need to enhance the depth and precision of organ-based classification is an essential step towards automation of medical operation and diagnostics. The research aims to investigate the effect and potential advantages transformer models have on binary semantic segmentation, the method utilized for the project. Hence, I employed the SegFormer model, for its lightweight architecture, as the primary deep learning model, alongside the Unet. A custom 2D computerized tomography (CT) scan dataset was assembled, CT-Org2D through meticulous operations. Extensive experiments showed that, in contrast to the selected models, the task’s simplicity required a redesigned Unet architecture with reduced complexity. This model yielded impressive results: Precision, Recall, and IOU scores of 0.91, 0.92, and 0.85 respectively. The research serves as a starting point, motivating further exploration, through different methodologies, to achieve even greater efficiency in organ segmentation.

## 1 Introduction

Fifteen million Americans have various operations performed annually, all of which carry a significant risk of unfavorable results. Several major issues may arise as a result of these operations’ decreased accuracy and precision. More painful scarring results from larger incisions made during surgery and from the surgeon’s limited range of motion to access certain areas. Apart from the potential for mistakes, these surgeries are very expensive overall, not only for the average patient but also for the hospitals that have to pay for the procedures.By 2025, hospital expenses are projected to average $40 million annually, accounting for medical staff, equipment, and other costs [2].

**Figure 1:**
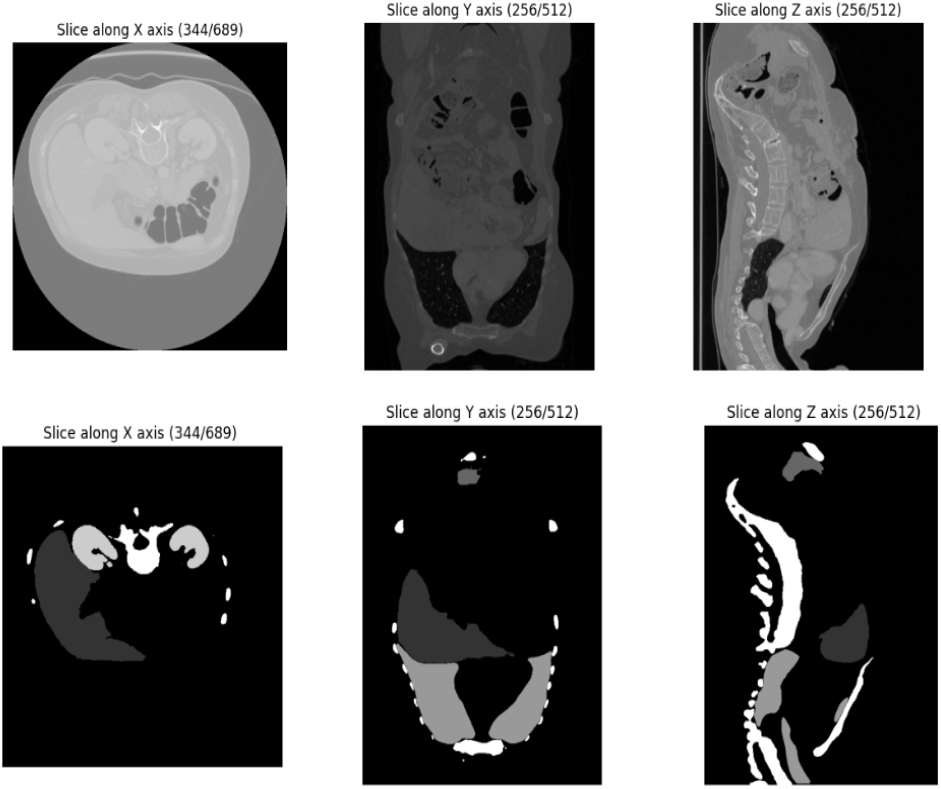
Different views in CT-Org’s volumetric 3D image with the features and their corresponding labels

With the implementation of robotic assistance, the necessity for human-medical assistance will grow obsolete as new advances in the intersection of the medical and AI worlds take place. The automation of surgical help has the potential to significantly change the medical industry by ensuring consistent precision, lowering surgical expenses, and enabling a reliable method of healthcare [9].

The initial stage in developing the knowledge required for automated procedures is through organ segmentation. This project emphasized binary semantic segmentation, where each pixel is labeled according to its class. The method uses downsampling for computational efficiency and upsampling to achieve the target image size, utilizing Max Pooling and Unpooling functions with specific strides and padding. Several architectures were explored and tested on, including transformers. Transformers have recently had a significant impact on the area of computer vision, which motivated me to research their limitations and abilities on semantic segmentation.

### 1.1 Model Architecture

Convolutional Neural Networks (CNN) are used to extract features, recognize patterns, and categorize various elements of an image. Traditional neural networks are slower and less effective at capturing features than CNNs. A CNN consists of: Convolutional, Pooling, and Fully-connected layers. All of the nodes are connected by these layers, which also help to extract features using filters and kernels by lowering complexity and parameter requirements. One of the characteristics that makes them successful is their ability to preserve information while reducing the high dimensionality of images.

**Figure 2:**
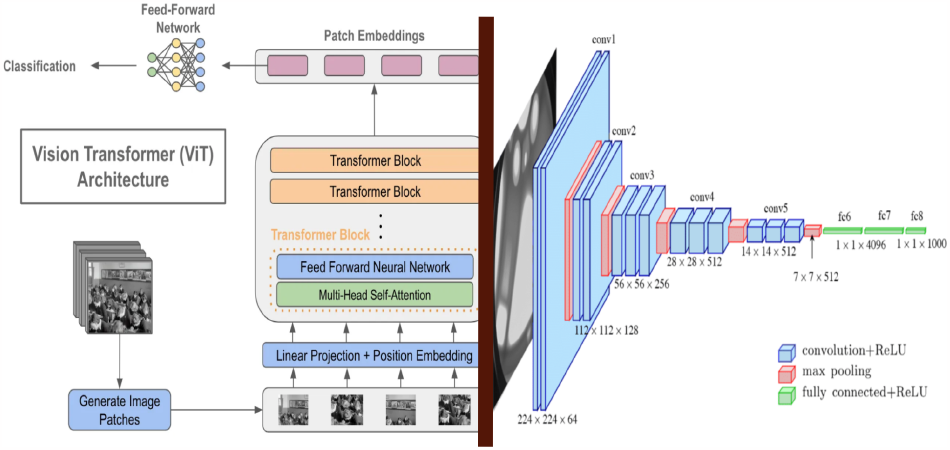
A visual representation of architectures from both Transformers(left) and CNNs(right)[2].

Parallel processing and self-attention mechanisms are features of the Transformers architecture, which is frequently employed in Natural Language Processing (NLP). Its selfattention component gives various components of an image different weights of importance. This is essential for capturing long-range dependencies, relationships between patches that are far apart from one another. The ability for independent and simultaneous evaluation of certain data points is known as parallel processing. In contrast, a recurrent neural network (RNN) examines each input point separately and dependently uses previous input for the current one. Transformers require significantly more data than the typical CNN, and as a result of this and parallel processing, their training times are typically longer.

## 2 Related Works

Numerous approaches and data types have been used in considerable research on the effectiveness of models in a variety of activities. Zettler et al. show that 2D U-Net models are more effective than 3D U-Net models in terms of speed and low memory costs [3]. Even while the 3D model had slightly more favorable results than the other model, they all concluded that this couldn’t possibly justify the computer resources that were wasted on it. The 3D model also worked with its respective 3D images data, which increased, even more, memory usage and rendering time. Similarly, Dia et al. use transformer-based architectures for multi-modal medical imaging classification [4]. The authors discuss their struggle to find a solution to the problem of medical imaging tasks not having sufficient data required to work with transformers. By creating a model that combines the benefits of a CNN’s low level feature extraction and a transformer’s long range dependencies, they are able to tackle this problem.

## 3 Methodology

I was motivated to select a 3D volume dataset (Ct-Org) with 6 classes: liver, lung, bladder, kidney, bone, and brain, to address the lack of 2D data online and customize it towards the task [1]. The data was gathered from CT scans of patients who had conditions or lesions in one or more of the organs specified. The images came from a wide variety of sources, including abdominal and full-body; contrast and non-contrast; and low-dose and high-dose CT scans. I condensed the original scans from 512 × 512 with varying depths, ranging from 74 to 987, to slices of 285 × 277 by rotating the view from axial(bottom view) to coronal(front view) for better organ visibility. This resulted in 512 slices per volume, creating a total of 71680 slices. The data was split into training (75%) and testing (25%) sets by volume to maintain slice grouping, and 20% of the training set was further separated for validation. The slices were sorted and identified using a specific naming convention(Ex.Volume0_Slice-0).

**Figure 3:**
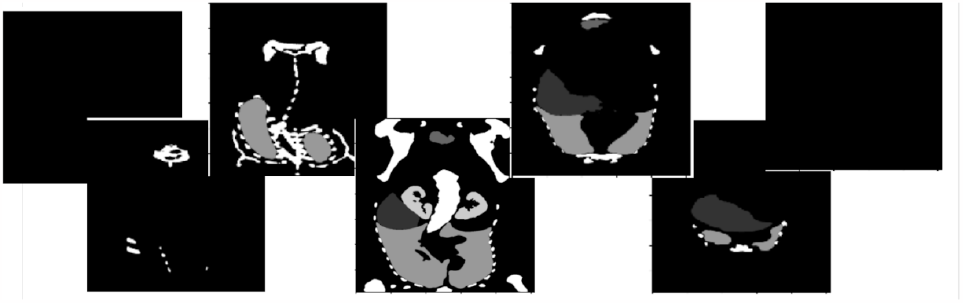
Sequential Slices at different depths/layers in a 3D labeled volume

### 3.1 Model Selection and Parameters

To represent the findings of transformers as a whole, the SegFormer model was chosen for this task, alongisde Unets, because of its functionality in semantic segmentation. This model avoids complex decoders, making the model lightweight. This results in the MLP decoders combining information from different layers [5]. Ultimately, its simple design contributes to its efficiency.

### 3.2 Data Preprocessing

In my new dataset, Ct-Org2D, the features (input) were grayscaled, but the labels were RGB. I duplicated the grayscaled channel twice so I could work with different models, then I combined all the outputs to get a three-channel image. This prevented the color from being added and also allowed me to use the 3-channel features and labels with ease. Alternatively, I may have customized the model’s layers to meet my singular channel requirement.

### 3.3 Model Training Process

For both the training and testing, the learning rate and batch size were set to 0.001 and 32, respectively. Both of which caused the speeds to become significantly slower. The Adam optimizer was chosen over SGD for its adaptiveness to adjust the weights better. The models ran for 20 epochs, but the scores usually converged after 8-10 epochs

## 4 Experiments

### 4.1 Experiment Setup

Google Colab Pro was used for running the code. The software gave access to efficient GPUs such as A100 and NVIDIA Tesla T4 for faster runtimes. Using higher memory settings, 25.5 GB of RAM was given. For the first half of the study, the PyTorch framework was employed for data collecting and analysis. Pytorch outperformed Keras and Tensorflow in terms of working-well with huge datasets and overall flexibility. Pytorch, however, demonstrated to be less incompatible with debugging, displaying images, and ability to handle complex models, especially transformers[8]. Hence, for the training, testing, and visualization phases, I utilized Keras/Tensorflow 2.13.

**Figure 4:**
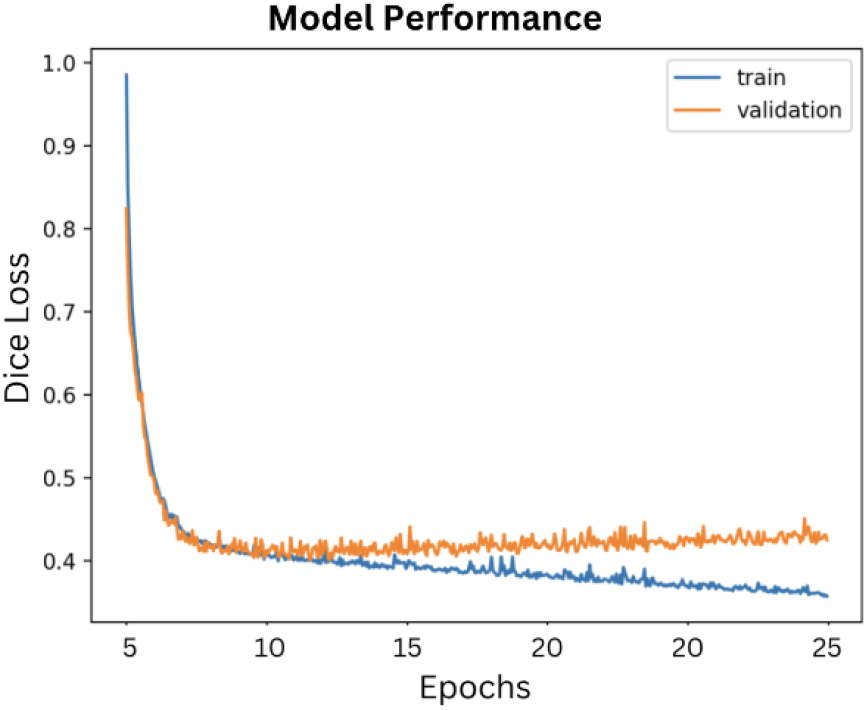
The graph depicts the convergence rates of a model across epochs using a Dice BCE Loss Function.

#### Important Libraries

1. SimpleITK - Extracts each slice and different characteristics(Ex. Shape, Views) from various volumes.
2. Matplotlib - Visualizes the predicted masks and input/output after being converted to a NumPy array.
3. Shutil - Relocates thousands of images to their respective training/testing drive folders.

### 4.2 Performace Metrics

To evaluate the performance, several key metrics were utilized: Precision, Recall, and IOU Score. Due to the task’s focus on binary segmentation, the IOU Score served as the benchmark metric for comparison. All metrics’ values range from 0 to 1, representing the probability in different aspects of the task: Each metric represents the probability, from 0 to 1, related to a unique element in the task:

#### Precision

The accuracy of true predictions relative to the entire set of outcomes[7]. In other words, it represents the proportion of events the model correctly predicted to be positive. To guarantee that non-organ regions are excluded from the segmentation, precision is essential.

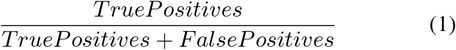

#### Recall

The measure of the percentage of actual positive results that the model accurately predicts. High recall is essential for segmenting organs since it guarantees thorough coverage of the entire organ. Insufficient recollection may result in insufficient segmentation, which may produce incorrect representations of the organ’s size and shape.

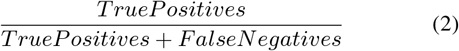

#### IOU Score

The overlap between the predicted regions and the ground truth. It is the most reliable metric for accurate binary segmentation.

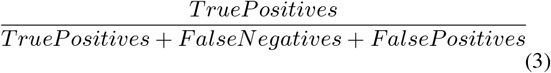

#### 4.3 Results

In Table 2, the evaluation scores are presented for precision, recall, and IOU score obtained from each model. It becomes evident that Segformer and the Unet models with more complex backbones had trouble adjusting to the binary segmentation task’s simplicity. This ultimately resulted in issues with overfitting. As a result, I developed a custom Unet model with fewer layers and nodes, which effectively reduced the size and complexity of the model as a whole. However, this model produced considerable results, supporting the idea that a model’s complexity must correspond to that of the task.

**Table 1:** Comparative data table of models. Showcases the different characteristics for all models employed in this study.

| Model | Parameters | Model Size | # of Layers |
| --- | --- | --- | --- |
| Custom-Unet | 550,000 | 2.2 mb | 51 |
| SegFormer | 47 million | 180 mb | 5 |
| UnetVgg16&19 | 31 million | 109 mb | 51 |
| UnetResnet50 | 20 million | 79.4 mb | 176 |

**Table 2:** Model performance metrics. Presents the results through various evaluation metrics, allowing for comparison in efficiency.

| Model | Precision | Recall | IOU Score |
| --- | --- | --- | --- |
| UnetVgg16 | 0.96 | 0.41 | 0.37 |
| UnetVgg19 | 0.95 | 0.35 | 0.36 |
| UnetResnet50 | 0.96 | 0.37 | 0.39 |
| Segformer | 0.56 | 0.33 | 0.26 |
| **Custom Unet** | **0.91** | **0.92** | **0.85** |

## 5 Conclusion

The study’s findings highlight a significant aspect of model selection and personalization. It becomes clear that for certain segmentation tasks, baseline models might not always yield the best results. Customization is a crucial tactic that enables alignment with the inherent features and challenges of the work. Notably, the performance of the custom Unet model, trained on a single dataset with minimal hypertuning and within a straightforward work environment showcases the potential for personalized model design. This demonstrates that even with minimal resources and adjustments, it is possible to achieve considerable improvements in accuracy and consistency.

Throughout the project, several limitations influenced the research process and results. One significant constraint was the computational resources limited by the work environment, including available RAM and software challenges. The size of the dataset made the problems much more difficult, leading to frequent timeouts and extended execution delays during debugging phases. Therefore, I created a more manageable subset of the data for troubleshooting.

### 5.1 Future Work

One could broaden the task’s scope to include multi-segmentation in order to improve this project even further. This would open the doors to more model options including intricate transformer architectures that would fit the complexity needs of the task. Another option is the addition of three dimensions to the task and utilizing various 3D datasets. The room for improvement is endless in the field of Organ Segmentation.

## Notes

### Competing Interest Statement

The authors have declared no competing interest.

